# Sociability genetically separable from social hierarchy in amniotes

**DOI:** 10.1101/2024.09.02.610763

**Authors:** Xin Lin, Guangyi Dai, Sumei Zhou, Yangyang Li, Yi-Hsuan Pan, Haipeng Li

**Affiliations:** CAS Key Laboratory of Computational Biology, Shanghai Institute of Nutrition and Health, University of Chinese Academy of Sciences, Chinese Academy of Sciences, 320 Yueyang Road, Shanghai, 200031, China; Key Laboratory of Brain Functional Genomics of Ministry of Education, School of Life Science, East China Normal University; Shanghai, China

**Keywords:** PAS1, tube test, social hierarchy, three-chamber social test, sociability, amniotes, social structure and organization

## Abstract

The emergence of social structure and organization is essential for the evolution of amniotes, including human beings. Sociability and social hierarchy are two key features to form a social group. However, it remains unknown whether sociability and social hierarchy are genetically separable. In this study, we examined the social hierarchy, social and social novelty preference of PAS1 (placental-accelerated sequence 1) knock-out and knock-in mice. PAS1 is a social enhancer that modulates social hierarchy. We found that PAS1 knock-out mice lack social hierarchy while wallaby/chicken PAS1 knock-in mice establish stable social ranks. Moreover, social and social novelty preference was observed in all PAS1-mutated mice. PAS1 knock-in mice have stronger preference to interact with other mice than wild-type mice (C57BL/6). No aggressive alteration was found in PAS1-mutated mice. Overall, our results showed that PAS1 is an indispensable regulatory element in the formation of social hierarchy while PAS1 regulates one of pathways modulating sociability. Therefore, sociability is genetically separable from social hierarchy in amniotes, providing insights into how social structure and organization evolved.

**Highlights:**

PAS1, a social enhancer, is indispensable for amniotes to establish social hierarchy.
PAS1 modulates sociability in amniotes.
Sociability and social hierarchy are regulated differently in amniotes.
The frequency of PAS1 allele was not rare when social hierarchy first appeared.

## Introduction

Sociability, the ability and tendency to have social interaction with others, is the basis and adaptive component of the social nature of many species, including human beings^1,2^. Social impairment and the lack of social interaction are considered the sign of multiple mental illnesses^3,4^. Another main characteristic of social animals is that the members of a group form different social levels, *i.e.*, social hierarchy^5,6^. To form social hierarchy, animals have to sense not only environmental changes^7–10^ but also the social status of each individual in the group^11,12^. In groups with stable hierarchical relationships, dominant animals have more resources than subordinate individuals, such as larger territory^13^, priority access to food, and more spouse options^14^. Rank in the hierarchical system greatly affects the behavior of animals, including mating, attack and stress responses^15^. Then a stable social structure and organization can be established, which is crucial to the survival of the group^16^. Therefore, the formation of social groups involves many developmental and neural factors, postnatal learning and memory.

It has been documented that the nonsocial-to-social and the social-to-nonsocial transition occurred multiple times during the evolution of amniotes^2,17^. Some amniotic species are solitary-living (nonsocial)^18^ while many amniotes are social^19^. The solitary-living indicates that an individual forages independently in its home range and encounters its mate only during the breeding season^20,21^. For those nonsocial or solitary-living species, the sociability is important for animals to seek their partners although those species lack social hierarchy^17^. Similarly, the identification of partners is also certainly crucial for social animals to maintain the social hierarchy and mate choices^22^. Therefore, sociability is an absolutely-necessary trait for all amniotes while social hierarchy is not. Since both of sociability and social hierarchy are coded in genomes, from an evolutionary perspective, sociability may be genetically separable from social hierarchy. However, this issue has never been investigated.

To address this important question, genetic components modulating sociability and social hierarchy should be studied. Since a nonsocial-to-social transition occurred in the ancestral lineage of placental mammals^17,23,24^, we recently developed a novel method to detect the accelerated evolution in both conserved and non-conserved genomic regions within the lineage^12^. Using this approach, we identified a social enhancer PAS1 (placental-accelerated sequence 1), located upstream of the *Lhx2* gene, a key transcription factor that regulates brain development^25,26^. Three PAS1 mouse strains with the C57BL/6 background were generated (PAS1*^-/-^*: PAS1 knock-out mice; PAS1*^w/w^*: wallaby PAS1 knock-in mice; PAS1*^c/c^*: chicken PAS1 knock-in mice). PAS1 knock-in mice (PAS1*^w/w^* and PAS1*^c/c^*) were able to maintain stable social ranks in cages with mixed homozygotes and heterozygotes of mutated mice while stable social ranks cannot be established in PAS1 knock-out mice^12^. Moreover, no structural abnormalities were found in the brain of PAS1-mutated mice. Overall, PAS1*^-/-^* is the first mouse strain identified to lack social hierarchy and social ranks can be turned over by mutating PAS1.

In this study, we examined the social rank stability and the sociability of the PAS1-mutated mice, focusing on the role of the enhancer PAS1 in modulating the two complex traits. Our findings provide strong evidences that PAS1 is an indispensable regulatory element in the formation of social hierarchy because PAS1 knock-out mice lack social hierarchy. We also found that PAS1 only regulates one of the pathways modulating sociability. Overall, these evidences indicate that sociability and social hierarchy are genetically separable in amniotes.

## Results

### Stable/unstable social ranks of PAS1-mutated mice

To examine the social hierarchy of PAS1-mutated mice, we detected the social rank of caged homozygote PAS1-mutated mice and wild-type mice, using the social dominance tube test^27,28^. Four male mice with the same genotype were housed together in each cage for at least eight weeks before the test (Table S1). It should be noted that we housed mice with the same genotype in one cage in this study. Differencing from our previous study^12^, where mice with different genotypes were housed in the same cage to examine whether PAS1-alleles modulate the social rank^12^. During the test, each mouse competed against its three cage mates to determine its social rank. When the ranks of four mice remained unchanged in one cage for at least four consecutive days, we considered their social ranks stable.

PAS1 knock-in and knock-out mice showed varying abilities to establish stable social ranks. We found that all caged PAS1*^w/w^* and PAS1*^c/c^* mice (wallaby and chicken PAS1 knock-ins) establish stable social ranks, similar to the patterns were observed in caged wild-type ones (Figure 1B, D, E). In contrast, caged PAS1*^-/-^* mice failed to establish stable social ranks (Figure 1C, and S1), even after nearly 20 weeks of housing. We also conducted social dominance tube test on PAS1 knock-out heterozygotes (PAS1*^m/-^*), all of which established stable social ranks (Figure S2A). There are not significant differences in time spent in home cage among different strains (Figure S3A, Table S1), nor were there any links between social rank and the body weight (Figure S3B). These evidences indicated that at least one PAS1-allele in each individual is essential to establish social hierarchy.

**Figure 1.**
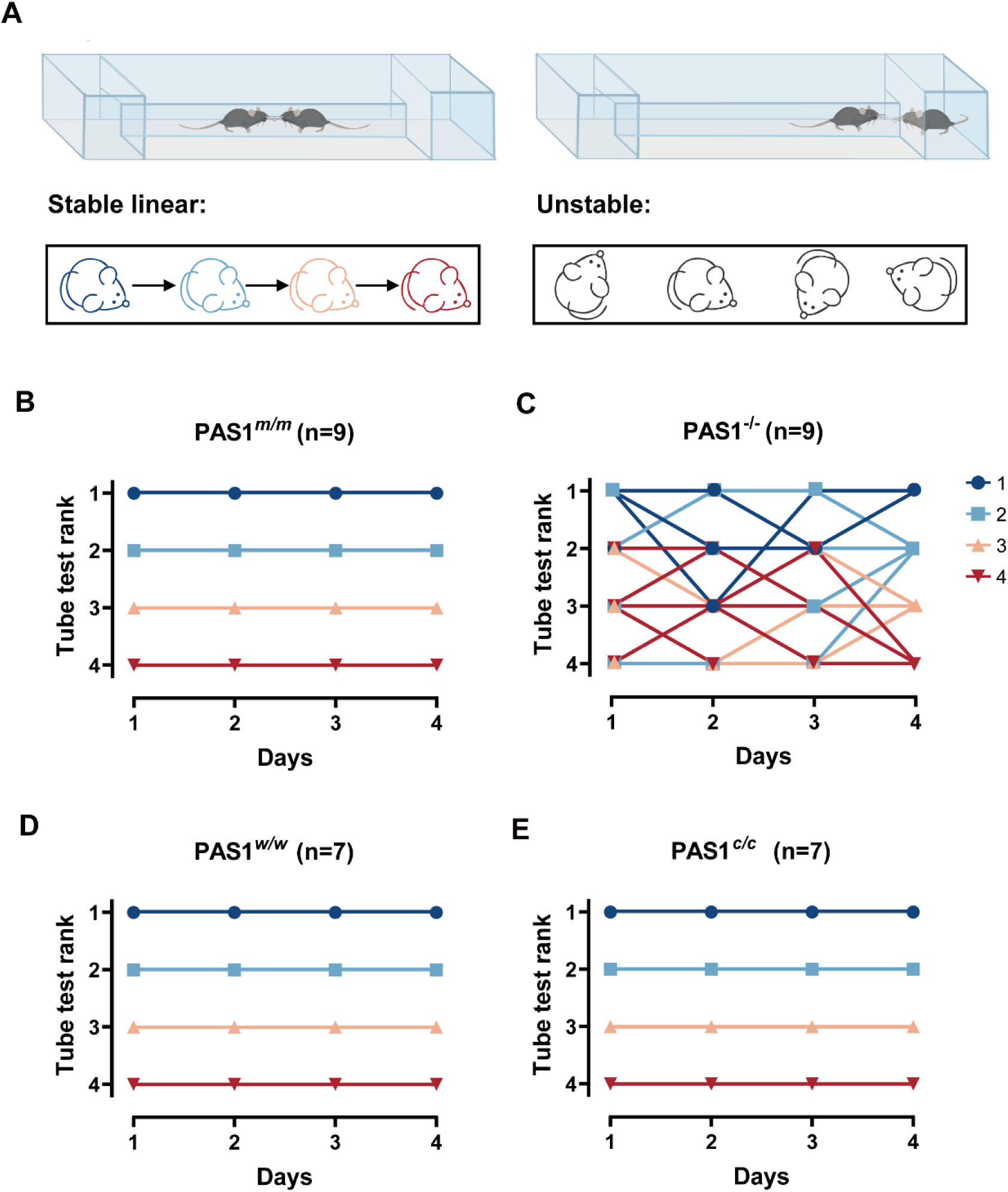
Stable/unstable social ranks of PAS1-mutated mice. **A.** Schematic diagrams of the social dominance tube test and social ranks of caged mice. Stable linear: four mice in one cage have stable linear ranks; Unstable: mice cannot gain stable ranks. **B-E.** Social ranks of PAS1-mutated homozygote mice. The colors are assigned according to the ranks obtained on the last day. PAS1*^m/m^*: wild-type mice; PAS1*^−/−^*: PAS1 knock-out mice; PAS1*^w/w^*: wallaby PAS1 knock-in mice; PAS1*^c/c^*: chicken PAS1 knock-in mice. *n*: number of cages tested.

### Sociability of PAS1-mutated mice

Since PAS1 knock-out homozygotes fail to form social hierarchy, it is important to examine the sociability of three PAS1-mutated strains. We investigated their social and social novelty preference using the three-chamber social test^29–32^. During the social preference stage, there was one circular wire cup in the corner of left chamber that held a juvenile stranger wild-type mouse (stranger1) (Figure 2A), while another empty cup was left in the diagonal corner of right chamber. The experimental wild-type mouse usually explores and sniffs around stranger1, demonstrating social preference (Figure 2C). We found that the total interaction time between experimental mice and stranger1 was very significantly higher than that between the experimental mice and empty cup (Paired *t*-test, PAS1*^m/m^*: *P* < 0.0001; PAS1*^-/-^*: *P* < 0.0001; PAS1*^w/w^*: *P* < 0.0001; PAS1*^c/c^*: *P* < 0.0001) (Figure 2E). Therefore, all the tested groups, including wild-type, PAS1 knock-out, wallaby PAS1 knock-in, and chicken PAS1 knock-in mice showed social preference.

**Figure 2.**
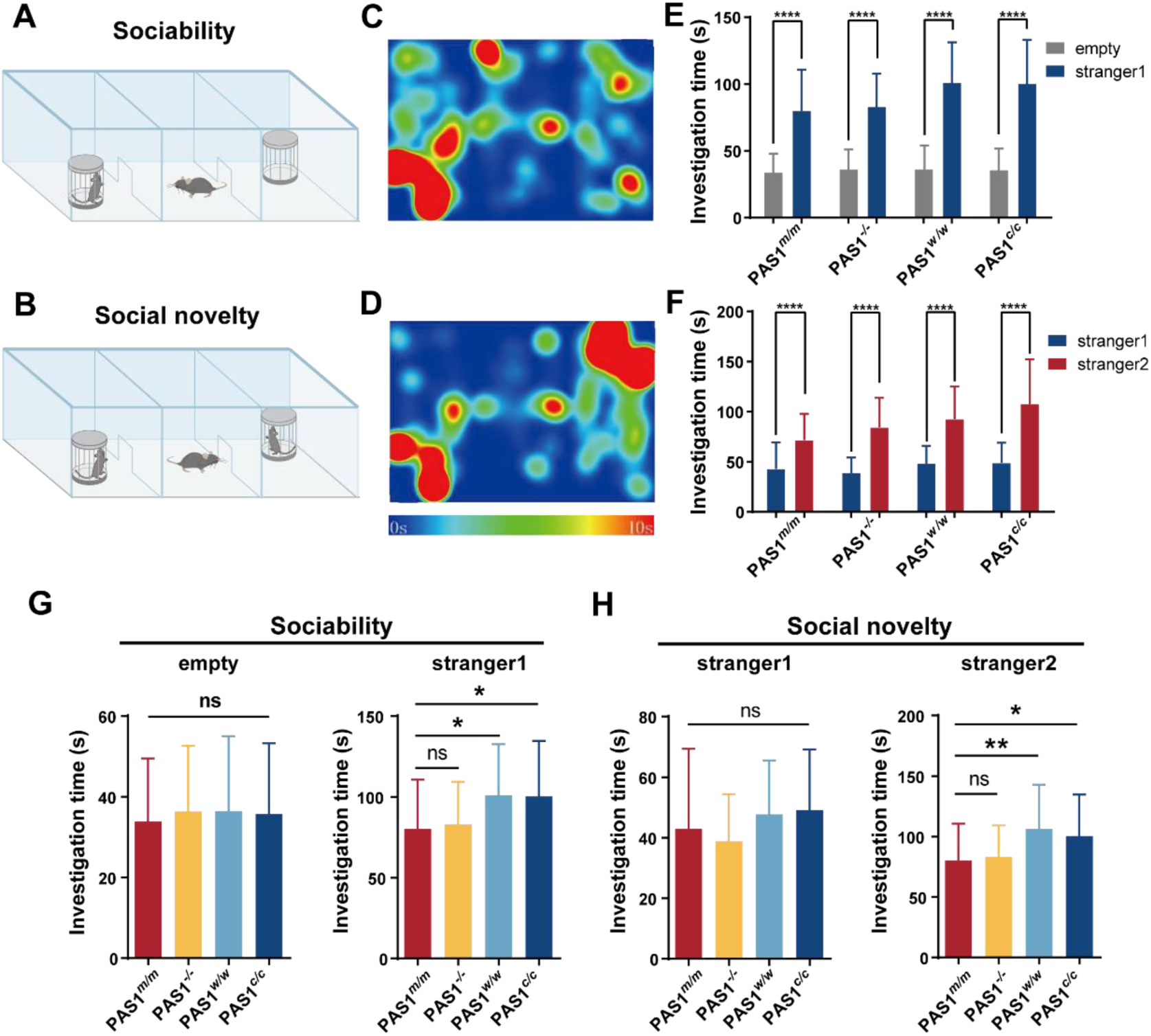
Sociability of PAS1-mutated mice. **A.** Schematic diagram of social preference stage in three-chamber social test. **B.** Schematic diagram of social novelty preference stage. **C-D**. Two examples of heatmap of mouse motion trail in the social and social novelty preference stages. **E.** Investigation time of experimental mice toward empty stimuli and stranger1 during social preference stage. **F.** Investigation time of experimental mice toward social familiar (stranger1) and un-familiar (stranger2) mice during social novelty preference stage. **G-H**. Investigation time of experimental mice toward social stimuli during the social and social novelty preference stages. Error bars represent standard deviation. PAS1*^m/m^*: n=36; PAS1^−/−^: n=36; PAS1*^w/w^*: n=33; PAS1*^c/c^*: n=33 individuals. \**P*<0.05, \*\**P*<0.01, \*\*\*\**P* < 0.0001, ns – not significant.

To examine whether PAS1 modulates the sociability, we compared the total social interaction time between three PAS1-mutated strains and the wild-type. Three PAS1-mutated strains, like the wild-type, showed no-interest in empty cup (Figure 2G). However, compared to the wild-type, both of two PAS1 knock-in strains showed a significantly greater interest in stranger1 (Ordinary one-way ANOVA, PAS1*^w/w^*: *P* = 0.015; PAS1*^c/c^*: *P* = 0.0201) (Figure 2G), demonstrating that PAS1 modulates sociability. Moreover, we observed no difference in the interacting time between experimental mice and stranger1 when examining the wild-type and PAS1 knock-out strain. Overall, these evidences indicated that there are multiple pathways modulating sociability.

During the social novelty preference stage, the mouse used in the social preference stage (stranger1) remained in the left chamber while another stranger wild-type mouse (stranger2) was held in the right chamber (Figure 2B). We found that all PAS1-mutated mice spend more time with stranger2 than with stranger1 (Paired *t*-test, PAS1*^m/m^*: *P* < 0.0001; PAS1*^-/-^*: *P* < 0.0001; PAS1*^w/w^*: *P* < 0.0001; PAS1*^c/c^*: *P* < 0.0001) (Figure 2D, F), indicating that PAS1-mutated mice have normal social memory and social novelty recognition. We then compared the interacting time between experimental mice and stranger1 and found that there is not significant difference between the wild-type and PAS1-mutated mice (Figure 2H). Similarly, when examining the interacting time between experimental mice and stranger2, there is not significant difference between the wild-type and PAS1 knock-out mice. However, the interacting time was significantly increased in both wallaby- and chicken-PAS1 knock-in mice, compared to the wild-type (Ordinary one-way ANOVA, PAS1*^-/-^*: *P* = 0.9597; PAS1*^w/w^*: *P* = 0.0027; PAS1*^c/c^*: *P* = 0.0274). We also examined PAS1 knock-out heterozygotes and found that PAS1*^m/-^* mice have normal social and social novelty preference (Paired *t*-test: *P* = 0.001 and 0.0085) (Figure S2B). Overall, these results suggested that PAS1 modulates social novelty preference.

### Dynamics of social and social novelty preference of PAS1-mutated mice

To further explore the dynamics of social and social novelty preference behavior in PAS1-mutated mice, an in-depth analysis of three-chamber social test results was performed. The sociability stage and the social novelty stage were partitioned into 5-second bins and the mean interaction time was obtained for each bin (Figure 3A). All experimental PAS1-mutated mice interacted more with unfamiliar stimulus (*i.e.*, stranger1 during the sociability stage and stranger2 during the social novelty stage) at the beginning and then their social interest gradually decreased over time. This pattern was similar to which was observed in the wild-type. During the early exploratory phase (first three minutes) (Figure S4A) of the sociability stage, the interaction time of experimental mice with stranger1 was much longer than to the empty cup. In the late exploratory phase (last three minutes) (Figure S4B), wild-type and PAS1 knock-out mice showed diminished interest in stranger1 while wallaby- and chicken-PAS1 knock-in mice remained a high level of interest. Similar observations were obtained during the social novelty stage. Overall, the wallaby- and chicken-PAS1 knock-in mice generally showed stronger interests in stranger mice than the wild-type mice, while all of them exhibited no interest in empty cup (Figure S4C).

**Figure 3.**
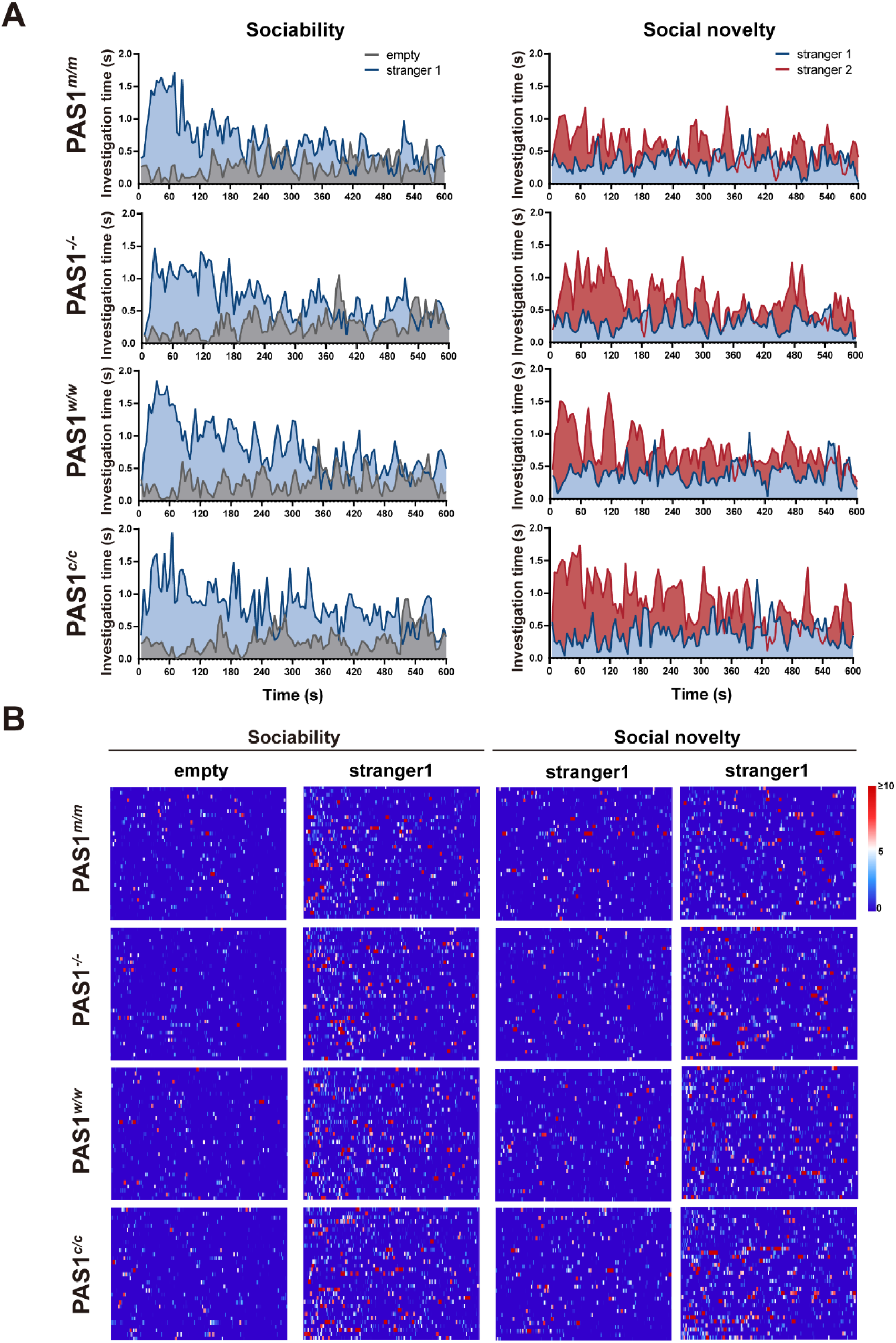
Dynamics of social and social novelty preference of PAS1-mutated mice. **A.** Investigation time of experimental mice, averaged in every five seconds, during social and social novelty preference stages. **B**. Heatmap of the investigation behavior of experimental mice in the first six minutes during social and social novelty preference stages. Each row represents one experimental mouse. PAS1*^m/m^*: n=36; PAS1*^−/−^*: n=36; PAS1*^w/w^*: n=33; PAS1*^c/c^*: n=33 individuals.

We then compared the dynamics of social and social novelty preference between PAS1-mutated mice and the wild-type. The investigation behavior of each experimental mouse during the first six minutes of both stages was presented as heatmaps, reflecting short and long bouts (Figure 3B). The long interactions shown as long bouts were largely observed between unfamiliar stimulus and experimental mice. Generally, PAS1*^-/-^* mice showed social interest in unfamiliar stimulus as the wild-type, and they rarely interacted with empty cup and familiar stimulus. Overall, the dynamics of social and social novelty preference support our findings that PAS1 knock-in mice have a stronger social and social novelty preference than both PAS1 knock-out mice and the wild-type.

Since social rank influences social behavior in mice^33^, we examined whether the social rank affects the social and social novelty preference (Figure S5). All of the mice were grouped by their ranks if the social hierarchy was established. Naturally, PAS1 knock-out mice were excluded from this analysis because they failed to establish the social hierarchy. The results showed that mice of different ranks exhibited similar behavior in sociability (Paired *t*-test for each rank: *P* < 0.0001) and social novelty preference (Paired *t*-test for each rank: *P* < 0.0001) (Figure S5A, B). Additionally, the investigation time to strangers has no significant differences among different ranks (Figure S5C, D). Therefore, social rank does not affect social and social novelty preference behavior.

### Aggressive behavior of PAS1-mutated mice

Aggression is a common behavior in social animals^34^. When a new social group is initially formed, social hierarchy is established through the process involving aggressive behaviors, such as chasing, attacking, pinning down, and aggressive grooming^35,36^, although C57BL/6 mice are generally less aggressive than other strains^37–39^. We then used the resident–intruder test to assess the aggressive behavior in PAS1-mutated mice (Figure 4A). For this test, an experimental mouse was first separated from its cage mates and housed individually in a new cage for at least 24 hours before the test. Then a juvenile intruder mouse was placed into the experimental mouse’s cage. To measure the level of aggressive behavior, we recorded the latency to the first active social contact of experimental mouse (as resident) and the total interaction time between experimental mouse and intruder mouse (Figure 4B, C).

**Figure 4.**
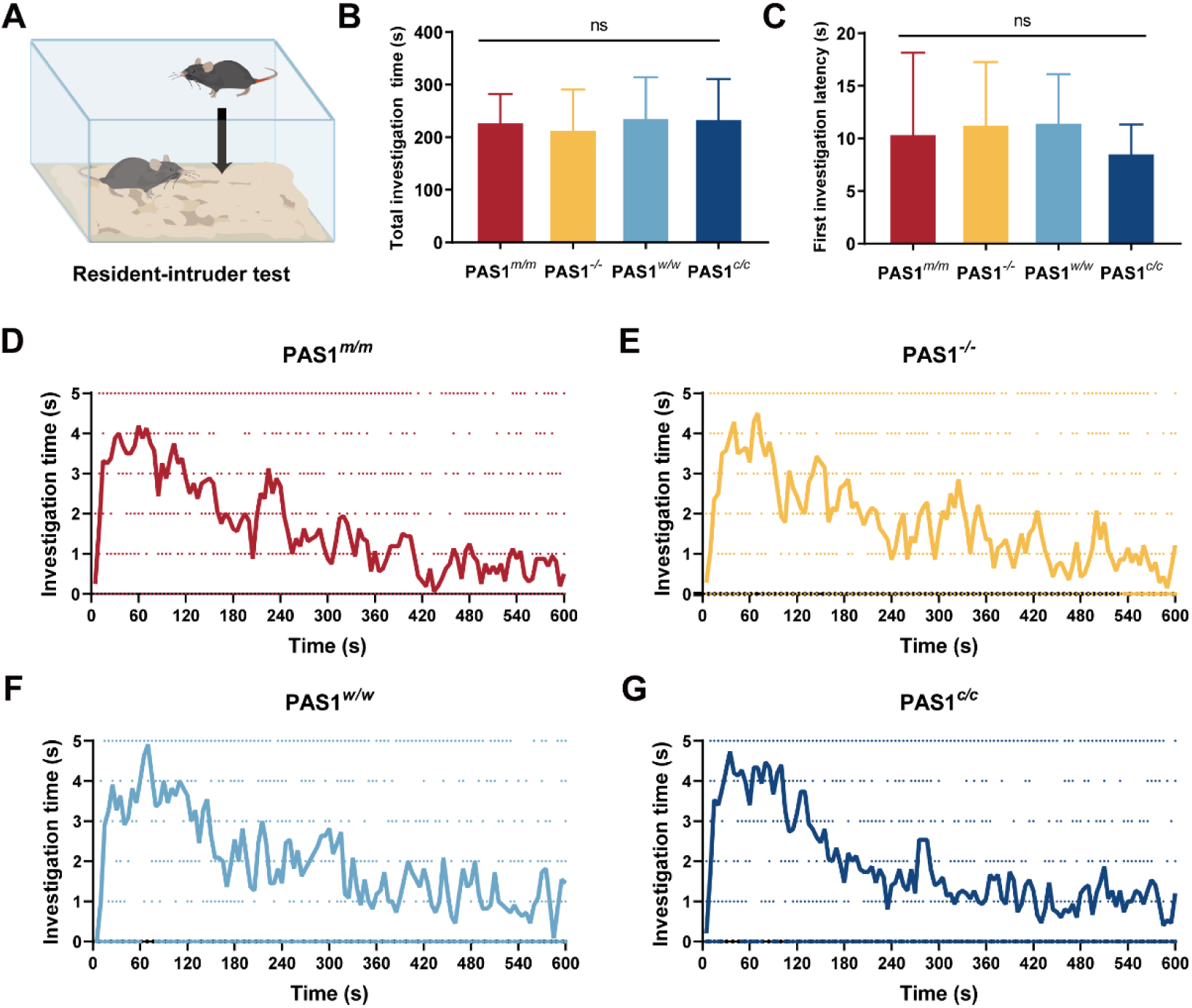
Aggressive behavior of PAS1-mutated mice. **A.** Schematic diagram of resident–intruder test. **B-C.** Total investigation time and the latency to the first contact time of PAS1-mutated mice during the resident–intruder test. **D-G.** Investigation time, averaged in every five seconds, during the resident– intruder test (PAS1*^m/m^*: n=13; PAS1*^−/−^*: n=14; PAS1*^w/w^*: n=11; PAS1*^c/c^*: n=14 individuals). Error bars represent standard deviation. ns – not significant.

All PAS1-mutated and the wild-type mice showed similar behavior to intruders in the test. The results of in-depth analysis demonstrated that the social interest of experimental mice in the intruders decreased over time in the wild-type and the three PAS1-mutated strains (Figure 4D-G). Thus, the mutated PAS1-alleles are unlikely to alter the aggressive behavior in mice and the lack of social hierarchy in PAS1 knock-out mice is not due to the aggression alteration.

## Discussion

Sociability and social hierarchy are strongly correlated, leading to the hypothesis that the regulatory mechanism of sociability and social hierarchy might be similar^34,40^. However, in this study we found that the social enhancer PAS1 plays a key role in the formation of social hierarchy while PAS1 regulates one of sociability-related pathways (Figure 5). Therefore, we suggested that sociability and social hierarchy are regulated differently in amniotes.

**Figure 5.**
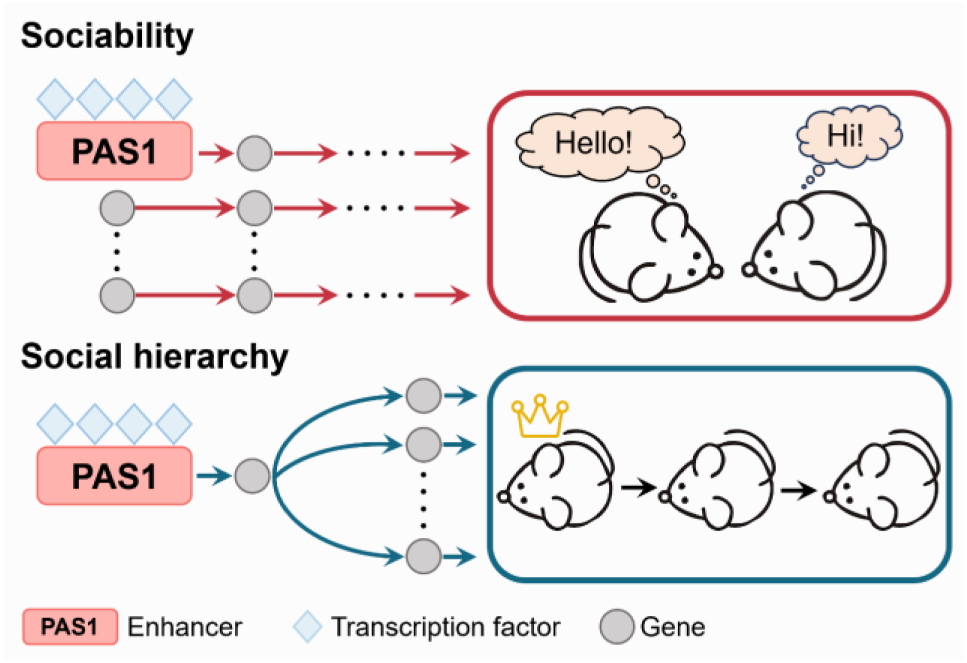
A model illustrating how sociability is genetically separable from social hierarchy in amniotes.

Since sociability is modulated by multiple pathways, genetic buffering and robustness is expected. This might be essential for the evolution of amniotes because genetic buffering and robustness ensures that the loss of function in a pathway, such as PAS1 knock-out, does not disable sociability. However, it remains to be further investigated how this redundancy has been maintained during the evolution although pleiotropy, the phenomenon of a single pathway influencing multiple traits, could be one of explanations.

It would be interesting to discuss how social hierarchy emerges after sociability occurs. We found that PAS1 knock-out heterozygotes (PAS1*^m/-^*) can establish stable social ranks (Figure S2A), demonstrating that a single copy of PAS1 allele is enough to cis-regulate downstream gene expression for the establishment of social hierarchy. However, social hierarchy cannot be established in cages with equally mixed PAS1 knock-out homozygotes and heterozygotes (PAS1*^-/-^* and PAS1*^m/-^*)^12^ because of the presence of PAS1*^-/-^*individuals. These findings indicate that, to establish social hierarchy in a population, each individual should have the ability to collaborate with others. This provides an important clue about how social hierarchy emerges in the nonsocial-to-social transition.

It is expected that a few individuals carrying a new PAS1 allele might have formed a group and established social hierarchy. The estimated lower bound of the frequency of the new PAS1 allele was 0.491 – 0.627 when social hierarchy first appeared in one group of an ancestral species by assuming that the group was composed of 10 – 20 diploid individuals (Figure S2C) (see the Method section). Therefore, the new PAS1 allele was not rare. We propose naming this phenomenon “population-dominant” because the phenotype of PAS1*^m/-^* relies on interactions with other individuals in the population, differing from classical dominance.

Different amniotic species, such as wolves, primates and meerkats, have diverse social structures and organizations. In many species, alpha males/females always play a key role in the social dynamics of those wild populations^41–44^. The replacement of dominant breeders is such an important event that affects the survival of these populations^45^. Therefore, we suggest that PAS1 and its downstream genes may have played roles in the determination of social structure and organization. When new beneficial mutations occurred on PAS1 and its downstream genes, these mutations should have got fixed quickly during the evolution of amniotes. These substitutions might alter the mechanisms to establish social hierarchy and subsequently affect the formation of social structure and organization. However, future works and large-scale experiments are needed to validate this hypothesis.

Overall, our study revealed that sociability is genetically separable from social hierarchy, which is essential for the evolution of amniotes. This finding evokes many important questions. Various brain regions have been confirmed to involve in the regulation of social behaviors in mice^20,27,46–48^, and it remains unclear in which brain regions and neural circuits PAS1 primarily acts. It is important to identify genes in PAS1-related pathways although a number of genes have been documented to influence social behaviors^49–51^. Additionally, it is essential to determine the developmental stages that are important for the functions of sociability and social hierarchy. These studies should help us to understand the nature of social structure / organization and provide new therapeutic target in mental-related diseases and stresses.

## STAR★Methods

### Key resources table

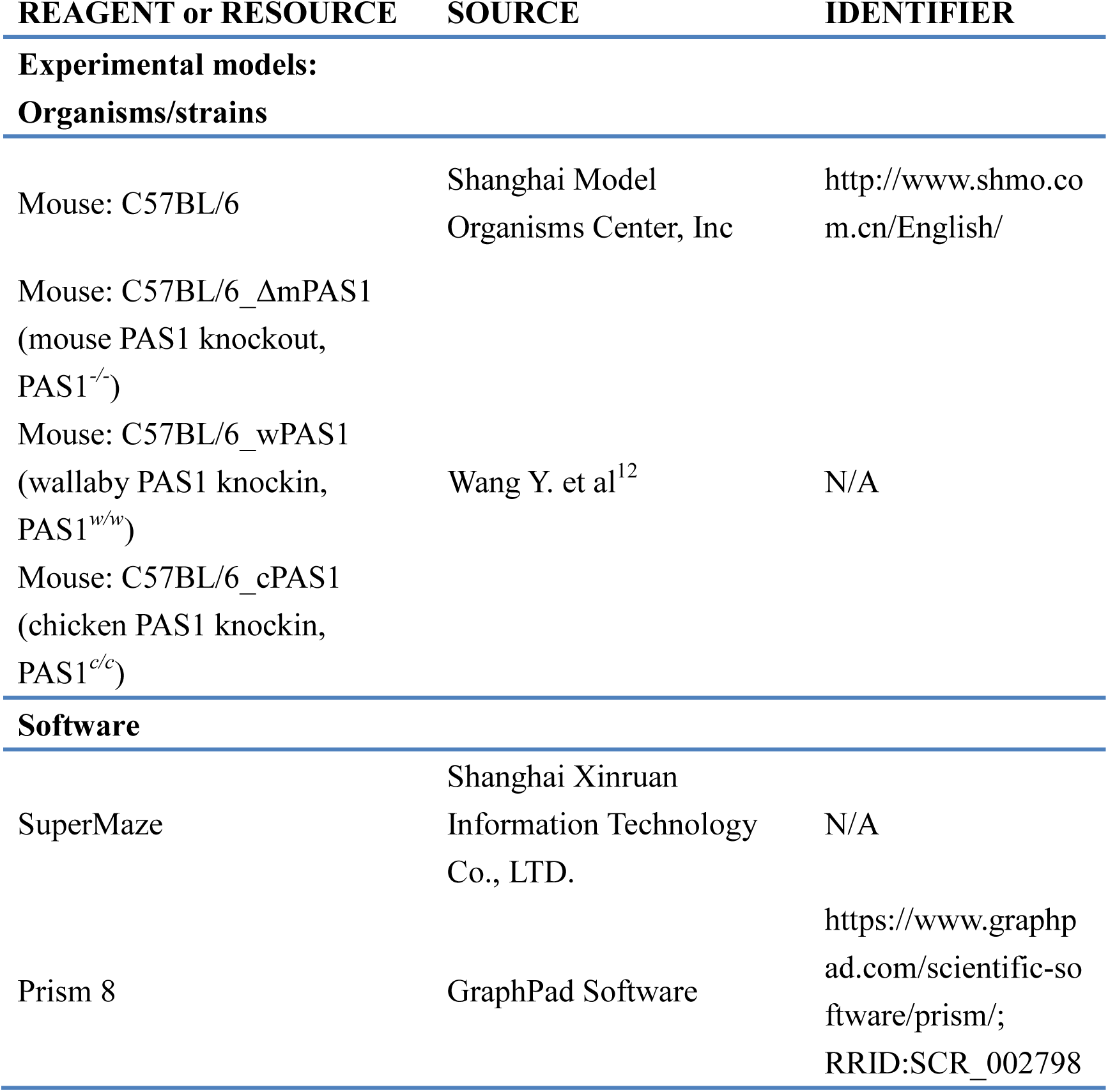

## Resource availability

### Lead contact

Further information and requests should be directed to and will be fulfilled by the lead contact, Haipeng Li.

## Materials availability

This study did not generate new unique reagents.

## Data and code availability

All data reported in this paper are enclosed in the supplementary materials. Any additional information should be available from the lead contact upon request.

## Experimental model and study participant details

### Animals

All animal experiments were performed in accordance with the protocols approved by the Committee and Laboratory Animal Department, Shanghai Institute of Nutrition and Health, Chinese Academy of Sciences. Governmental and institutional ethical guidelines were followed. PAS1 knock-out and knock-ins were generated in our previous study^12^. Generally, 9–24 weeks old male mice were used for testing behaviors. Mice were housed in specific-pathogen-free condition.

### Social dominance tube test

Four male mice of the same genotype with similar age and weight were cage-housed together for at least two weeks before the test. The age difference of these mice in one cage was less than 10 days. During the test, mice were allowed to run through a transparent plexiglass tube of 40 cm in length and 3 cm in diameter, a size just sufficient to permit one adult mouse to pass through without reversing the direction. There is a small chamber (17 ×8 ×14 cm) at each end of the tube for temporary housing purpose.

To prepare for the test, each experimental mouse was given eight training runs every day for three days. Immediately before the test, experimental mice were given four additional training runs. During the test, two male mice were released at each end of the tube. There is a moveable transparent plexiglass door in the middle of the tube to ensure that the two mice meet in the middle of the tube. If a mouse retreated from the tube within 2 min, it was given a score of zero. Another mouse that did not retreat was given a score of one. If no mice retreated within 2 min, the test was repeated. During the training stage, if no mice retreated in three successive trials, the test was carried out in the next day. These cases were rare and only observed in the early training stage of PAS1*^-/-^* mice. Between trials, the tube was cleaned with 75% ethanol. From trial to trial, experimental mice were released at either end alternatively. For each pair of male mice, two or three trials were conducted, and the mouse that won two times was considered the winner of the test. A new tube was employed for the next pair of mice. Mice were allowed to rest for at least 10 min between tests. Each mouse competed with its three cage-mates. To determine the stability of social ranks over time, mice were tested under the same conditions every day for at least seven days. The rank of an experimental mouse was assessed by the number of wins against its cage mates at the last day if the ranks remained stable for at least four consecutive days.

### Three-chamber social test

Male mice (between18-24 weeks old) were housed together according to standards and requirements for at least two weeks before the test^32^. The hand-made box (60×40×22cm) was divided into three chambers by clear plexiglass sheets with opening “door” to allow experimental mice to move between the chambers. A suitable white plexiglass sheet was used as the floor. Two identical circular wire cup-like containers, that are large enough to hold a single mouse, were used to hold stranger-mice. One was placed at the corner of the left chamber; the other was placed at the diagonal corner of the right chamber. The containers are comprised of metal wires to allow for air exchange and to prevent direct physical interactions between the inside animal (stranger) with the outside one (experimental mouse). In this study, 12-16 weeks old wild-type male mice were used as strangers.

Before performing the test, a habituation phase was done. Experimental mouse was placed into a clean three-chamber social approach box with two empty cups and allowed to explore the box freely for 10 minutes. For the sociability stage, experimental mouse was first placed in the center chamber while the doors between chambers were closed. A juvenile stranger mouse (stranger1) was put in the wire cup of the left chamber, while the right cup was empty (Figure 2A). And then the doors between chambers were opened to allow experimental mouse free access to explore three chambers for 10 minutes. For the sociability stage, a novel unfamiliar mouse was put in the previously empty cup of the right chamber, while the same familiar mouse used in the sociability stage (stranger1) was still in the left chamber (Figure 2B). Then experimental mouse was allowed to free explore three chambers for another 10 minutes. After each trial, all chambers and wire cups were cleaned with 75% ethanol and air-dried for 10 minutes, to prevent olfactory cue bias and to ensure proper disinfection. Social behaviors were recorded and analyzed by using SuperMaze (Shanghai Xinruan Information Technology Co., LTD.).

### Resident–intruder test

The day before the experiment, an experimental mouse (12-15 weeks old) was separated from its cage mates and lived alone for one day. On the next day, the experimental mouse was placed in a testing room with its home cage for 15 minutes. During the active (dark) phase of the light/dark cycle, a marked intruder was placed into the experimental mouse’s home cage. Intruders (8-10 weeks old) with dark coats were identified with nontoxic red paint at the base of their tails. Repeated use of the same intruders was allowed, but at least 2h between uses was required. Mice were then left undisturbed and they were videotaped freely for 10 mins. The measures for the resident–intruder test of the latency to the first active social contact of resident and the total interaction time between resident and intruder mouse.

### Statistical analyses

Statistical analyses were performed using Prism 8 (GraphPad Software). Values are presented as mean ±SD. All data were first tested with Shapiro-Wilk normality test for the normal distribution. To compare behaviors between two groups in three-chamber social test, *P-*values were calculated using the Paired *t*-test. For comparison of four groups with single independent variable in the resident-intruder test and three-chamber social test, *P*-values were calculated using the Ordinary one-way ANOVA followed by the post hoc test with Dunnett’s multiple comparisons. To analyze the single variable associated with the four ranks, *P*-values were calculated with RM one-way ANVOA followed by the post hoc test with Tukey’s multiple comparisons. \**P*<0.05, \*\**P*<0.01, \*\*\**P* < 0.001, and \*\*\*\**P* < 0.0001.

### Lower bound of PAS1 allele frequency to establish social hierarchy

Denote the ancestral non-functional allele of PAS1 by “*A*”, and the new PAS1 functional allele by “*a*”. Their frequency was *f*(*A*) = *p* and *f*(*a*) = *q*. We assumed that the “*A*” allele, such as the PAS1-null allele, has no function to establish social hierarchy. There were three genotypes in the ancestral species: *AA*, *Aa*, and *aa*. Their frequency was *p*^2^, 2*pq*, and *q*^2^, respectively. To establish social hierarchy, the genotypes of individuals in a group had to be *Aa* or *aa*. Therefore, the probability to form such a group is (1 − *p*^2^)*^n^*, by considering random sampling effects, where *n* is the number of individuals in the group. If we assumed *n* = 20 and required that this probability is larger than 0.05, we obtained *f*(*AA*) ≤ 0.139 (Figure S2C), indicating that the lower bound of frequency of new PAS1 functional allele *f*(*a*) is 0.627. We also obtained the lower bound of *f*(*a*) to be 0.491 when *n* = 10. Overall, these results indicated that the frequency of new PAS1 functional allele should not be low when social hierarchy first established in a group during a nonsocial-to-social transition.

## Acknowledgements

We thank Dr. Ben Zhou and the animal caretaker team for assistance with animal care and experiments. This work was supported by the National Natural Science Foundation of China and the Chinese Academy of Sciences.

## Author Contributions

XL, GD, SZ, YL, YHP and HL conceived the study. XL, GD, SZ and YL collected and analyzed the data. All authors discussed the results and wrote the paper.

---

**Figure S1.**
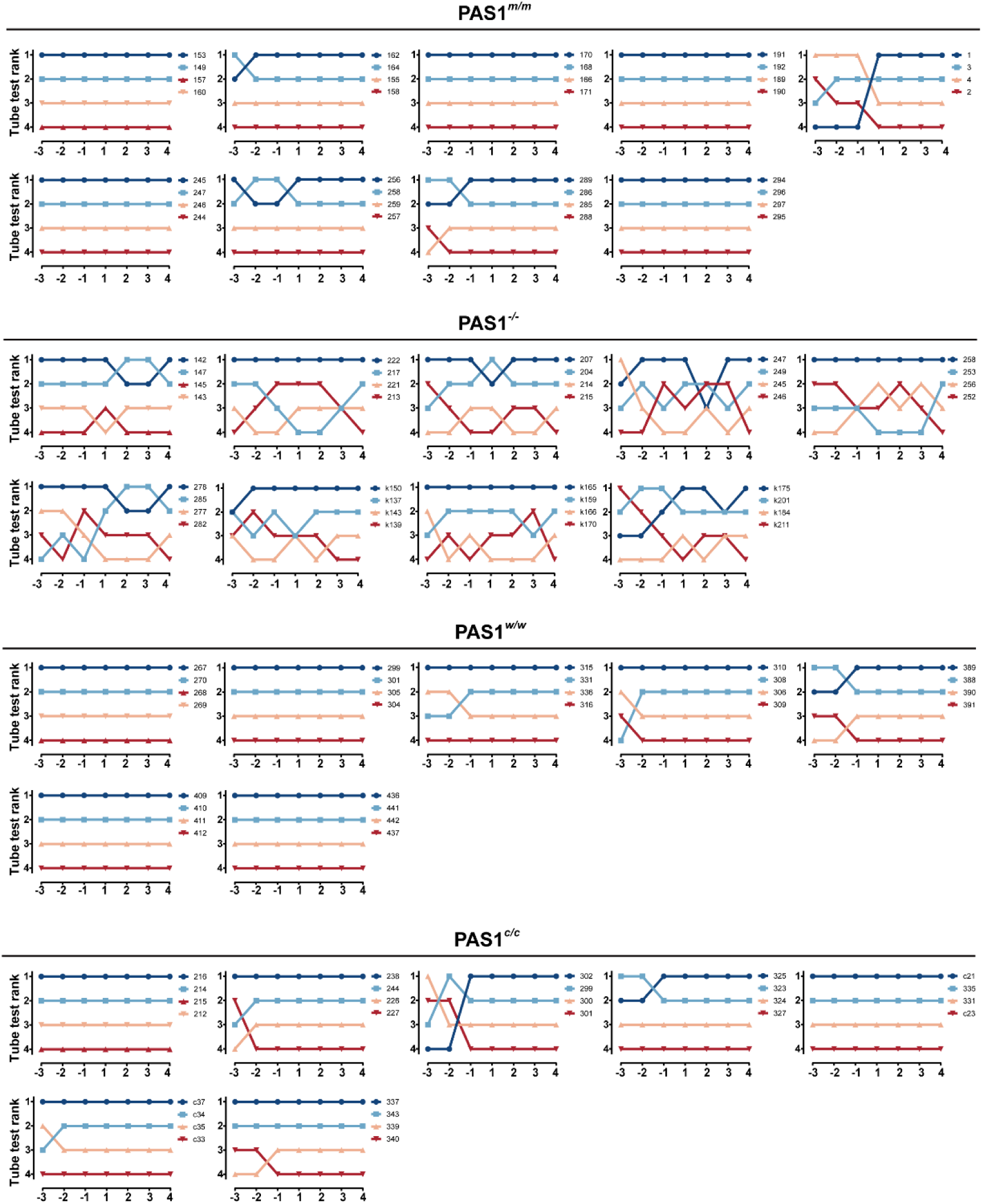
Results of social dominance tube test for each examined cage. Tube-test ranks of mice in each cage in seven consecutive days. Days numbered by negative represent the training days.

**Figure S2.**
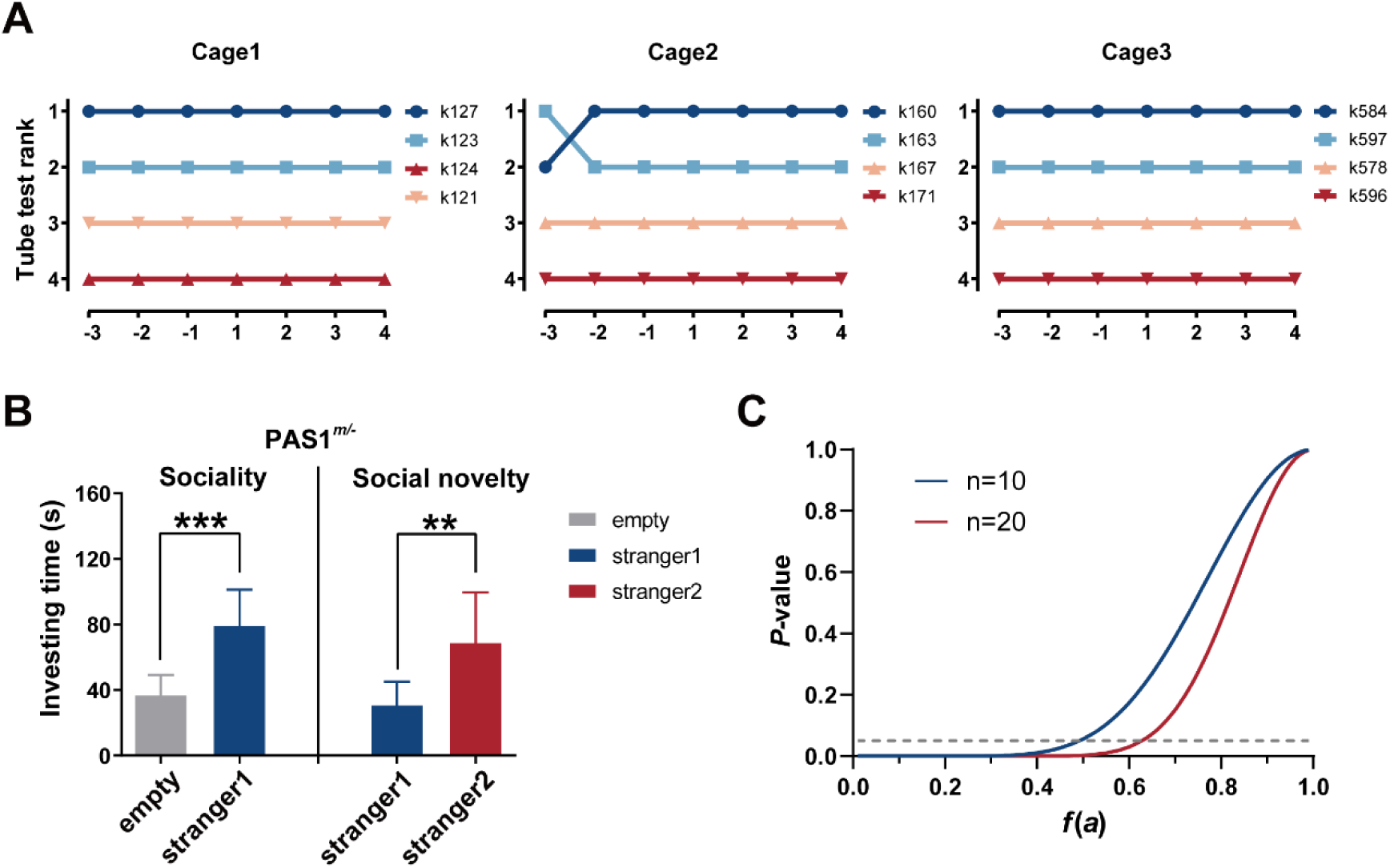
Social behaviors of PAS1 knock-out heterozygotes (PAS1*^m/-^*) and population-dominant phenomenon. **A.** Tube-test ranks of PAS1 knock-out heterozygous mice (PAS1*^m/-^*) in three cages in seven consecutive days. Days numbered by negative are the training days. **B.** Investigation time of experimental mice (PAS1*^m/-^*) toward social stimuli during the three-chamber social test. \*\**P* < 0.01, \*\*\**P* < 0.001. Error bars represent standard deviation. **C.** The probability to form a group with the genotypes of *Aa* or *aa* (the *y*-axis), given the frequency of *a* in the ancestral species (the *x*-axis). The “*A*” allele denotes the ancestral non-functional allele of PAS1, and the “*a*” allele is the new PAS1 functional allele. The dotted line indicates that the probability is 0.05. *n* is the number of individuals in the group.

**Figure S3.**
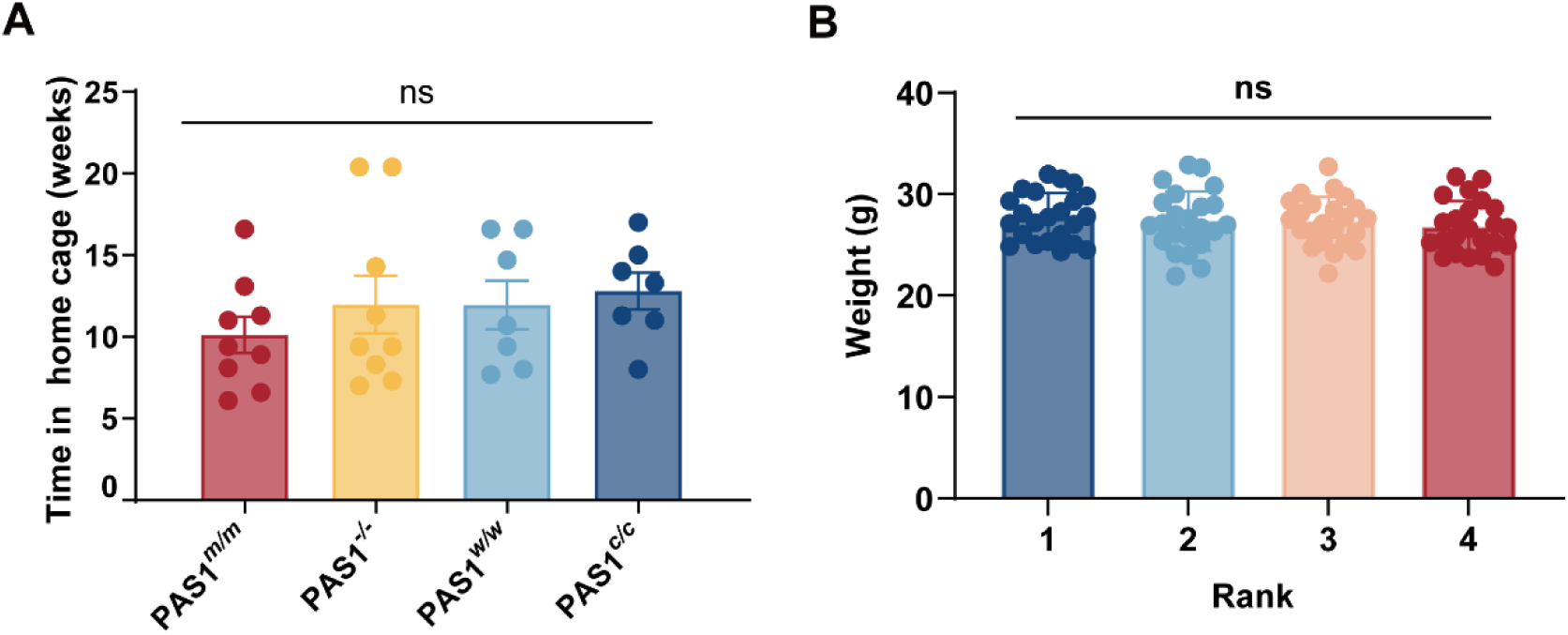
Basic parameters of mice in social dominance tube test. **A.** Time in home cage. PAS1*^m/m^*: *n* = 9 cages; PAS1*^-/-^*: *n* = 9 cages; PAS1*^w/w^*: *n* = 7 cages; PAS1*^c/c^*: *n* = 7 cages. **B.** Weight of mice categorized according to their social ranks. *n* = 23 individuals (for each rank). Error bars represent standard deviation, ns – not significant.

**Figure S4.**
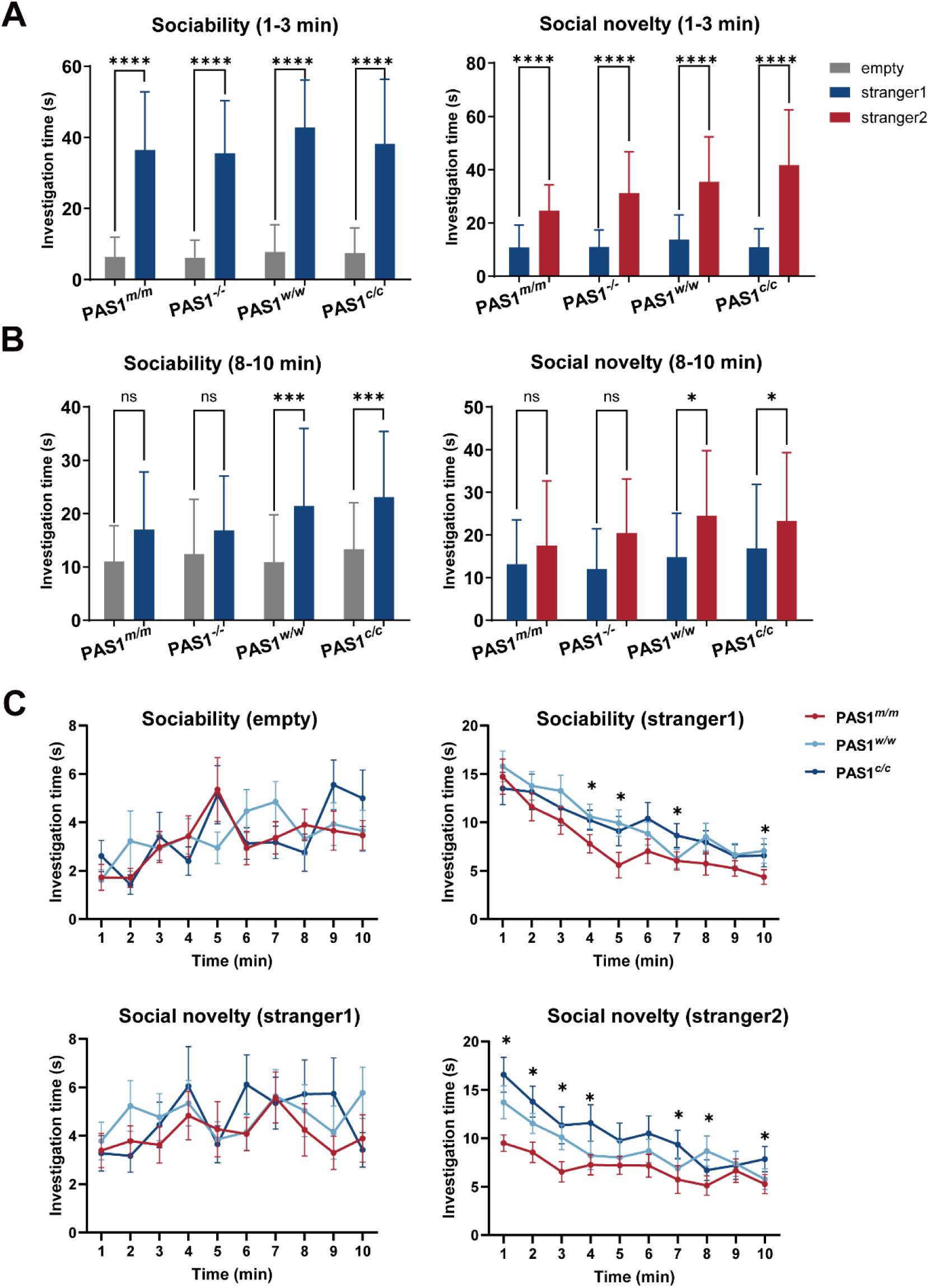
Comparisons of PAS1-mutated mice in social and social novelty preference stages. **A.** Investigation time of experimental mice in the first three minutes of social and social novelty preference stages. **B.** Investigation time of experimental mice in the last three minutes of social and social novelty preference stages. **C.** Investigation time of experimental mice through social and social novelty preference stages. PAS1*^m/m^*: n=36; PAS1^-/-^: n=36; PAS1*^w/w^*: n=33; PAS1*^c/c^*: n=33 individuals. Error bars represent standard deviation. \**P* < 0.05, \*\**P* < 0.01, \*\*\**P* < 0.001, \*\*\*\**P* < 0.0001, ns – not significant.

**Figure S5.**
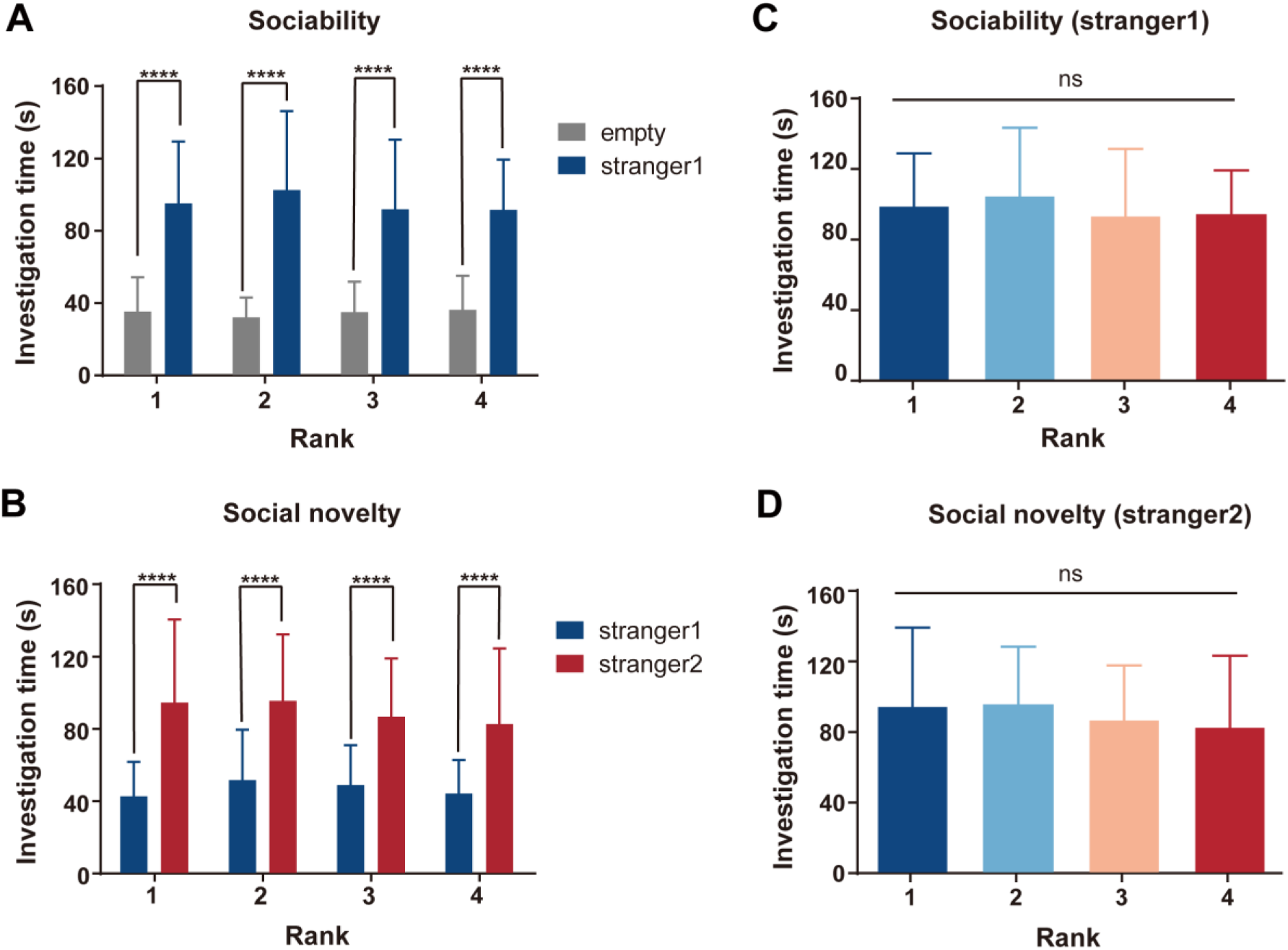
Sociability of mice categorized according to their social ranks. The results of PAS1 knock-out mice were discarded here because these mice failed to establish stable social ranks. **A.** Investigation time of experimental mice toward stranger1 and empty stimuli during the social preference stage. **B.** Investigation time of experimental mice toward familiar (stranger1) and un-familiar (stranger2) mice during the social novelty preference stage. **C.** Investigation time of experimental mice toward stranger1 during the social preference stage. **D.** Investigation time of experimental mice toward stranger2 during the social novelty preference stage. \*\*\*\**P* < 0.0001; ns – not significant. Error bars represent standard deviation.

**Table S1.** Basic parameters of mice in social dominance tube test.

| Genotype | Cage number | Age range at group-housing (weeks) | Age range on tube-test day (weeks) | Weight range on tube-test day (g) |
| --- | --- | --- | --- | --- |
| PAS1<sup>m/m</sup> | Cage 1 | 3.1 | 12.6 | 24.3-25.8 |
|  | Cage 2 | 3-3.7 | 11.9-12.6 | 24.1-29.3 |
|  | Cage 3 | 3.1-4.6 | 19.7-21.1 | 27.7-31.5 |
|  | Cage 4 | 3.1 | 14.1 | 22.7-26.6 |
|  | Cage 5 | 3.0 | 11.1 | 23.7-28.3 |
|  | Cage 6 | 2.9 | 9.4 | 27.0-28.5 |
|  | Cage 7 | 2.7 | 8.9 | 23.8-25.1 |
|  | Cage 8 | 3.1 | 16.3 | 26.0-30.8 |
|  | Cage 9 | 3.1 | 14.4 | 26.7-30.5 |
| PAS1<sup>-/-</sup> | Cage 1 | 3.0 | 17.8 | 26.9-29.8 |
|  | Cage 2 | 3.0-3.6 | 23.4-24.0 | 28.4-33.3 |
|  | Cage 3 | 3.0 | 23.4 | 25.1-28.7 |
|  | Cage 4 | 3.3 | 17.6 | 24.1-29.1 |
|  | Cage 5 | 3.1-4.4 | 12.5-13.9 | 25.25-29.1 |
|  | Cage 6 | 10.9-11.9 | 17.9-18.9 | 26.1-30.0 |
|  | Cage 7 | 3.0-4.0 | 11.3-12.3 | 24.3-25.4 |
|  | Cage 8 | 3.0 | 10.3 | 24.1-28.0 |
|  | Cage 9 | 4.1-4.6 | 18.0-18.4 | 27.11-30.78 |
| PAS1<sup>w/w</sup> | Cage 1 | 3.0 | 17.7 | 25.4-30.0 |
|  | Cage 2 | 3.6 | 20.1 | 26.6-32.0 |
|  | Cage 3 | 3.1-3.6 | 12.8-13.0 | 22.8-26.2 |
|  | Cage 4 | 3.1 | 19.7 | 25.3-27.0 |
|  | Cage 5 | 2.7 | 10.7 | 24.9-28.1 |
|  | Cage 6 | 2.6 | 13.3 | 27.2-32.6 |
|  | Cage 7 | 3.0-3.6 | 10.7-11.3 | 27.2-29.8 |
| PAS1<sup>c/c</sup> | Cage 1 | 3.1-4.3 | 20.1-21.3 | 27.6-31.1 |
|  | Cage 2 | 3.0 | 18.0 | 25.2-27 |
|  | Cage 3 | 3.4 | 11.4 | 21.9-25.2 |
|  | Cage 4 | 3.0 | 14.3 | 27.5-29.3 |
|  | Cage 5 | 3.0 | 12.0 | 27.1-30.1 |
|  | Cage 6 | 3.0-3.3 | 17.0-17.3 | 30.3-32.8 |
|  | Cage 7 | 3.3 | 16.6 | 25.1-29.0 |

## References

1. Clutton-Brock, T. (2021). Social evolution in mammals. Science 373, eabc9699.

2. Wilson, E.O. (2000). Sociobiology: the new synthesis, twenty-fifth anniversary edition (Harvard University Press).

3. Toth, I., and Neumann, I.D. (2013). Animal models of social avoidance and social fear. Cell Tissue Res 354, 107–118.

4. Snyder-Mackler, N., Burger, J.R., Gaydosh, L., Belsky, D.W., Noppert, G.A., Campos, F.A., Bartolomucci, A., Yang, Y.C., Aiello, A.E., O’Rand, A., et al. (2020). Social determinants of health and survival in humans and other animals. Science 368, eaax9553.

5. Koski, J.E., Xie, H., and Olson, I.R. (2015). Understanding social hierarchies: the neural and psychological foundations of status perception. Soc Neurosci 10, 527–550.

6. Park, M.J., Seo, B.A., Lee, B., Shin, H.S., and Kang, M.G. (2018). Stress-induced changes in social dominance are scaled by AMPA-type glutamate receptor phosphorylation in the medial prefrontal cortex. Sci Rep 8, 15008.

7. Lihoreau, M., Charleston, M.A., Senior, A.M., Clissold, F.J., Raubenheimer, D., Simpson, S.J., and Buhl, J. (2017). Collective foraging in spatially complex nutritional environments. Philos Trans R Soc Lond B Biol Sci 372, 20160238.

8. Hannibal, D.L., Bliss-Moreau, E., Vandeleest, J., McCowan, B., and Capitanio, J. (2017). Laboratory rhesus macaque social housing and social changes: implications for research. Am J Primatol 79, 1–14.

9. He, K., Liu, Q., Xu, D.M., Qi, F.Y., Bai, J., He, S.W., Chen, P., Zhou, X., Cai, W.Z., Chen, Z.Z., et al. (2021). Echolocation in soft-furred tree mice. Science 372, eaay1513.

10. Li, Y.Y., Lv, Q.Y., Zheng, G.T., Liu, D., Ma, J., He, G.M., Zhang, L.B., Zheng, S., Li, H.P., and Pan, Y.H. (2022). Unexpected expression of heat-activated transient receptor potential (TRP) channels in winter torpid bats and cold-activated TRP channels in summer active bats. Zool Res 43, 52–63.

11. Li, Q., Wang, M., Zhang, P., Liu, Y., Guo, Q., Zhu, Y., Wen, T., Dai, X., Zhang, X., Nagel, M., et al. (2022). A single-cell transcriptomic atlas tracking the neural basis of division of labour in an ant superorganism. Nat Ecol Evol 6, 1191–1204.

12. Wang, Y., Dai, G., Gu, Z., Liu, G., Tang, K., Pan, Y.H., Chen, Y., Lin, X., Wu, N., Chen, H., et al. (2020). Accelerated evolution of an *Lhx2* enhancer shapes mammalian social hierarchies. Cell Res 30, 408–420.

13. Jarrell, H., Hoffman, J.B., Kaplan, J.R., Berga, S., Kinkead, B., and Wilson, M.E. (2008). Polymorphisms in the serotonin reuptake transporter gene modify the consequences of social status on metabolic health in female rhesus monkeys. Physiol Behav 93, 807–819.

14. Chou, Y.J., Ma, Y.K., Lu, Y.H., King, J.T., Tasi, W.S., Yang, S.B., and Kuo, T.H. (2022). Potential cross-species correlations in social hierarchy and memory between mice and young children. Commun Biol 5, 230.

15. D’Amato, F.R., Rizzi, R., and Moles, A. (2001). A model of social stress in dominant mice: effects on sociosexual behaviour. Physiol Behav 73, 421–426.

16. Zhou, T., Sandi, C., and Hu, H. (2018). Advances in understanding neural mechanisms of social dominance. Curr Opin Neurobiol 49, 99–107.

17. Lukas, D., and Clutton-Brock, T.H. (2013). The evolution of social monogamy in mammals. Science 341, 526–530.

18. Hansen, J.E., Hertel, A.G., Frank, S.C., Kindberg, J., and Zedrosser, A. (2022). Social environment shapes female settlement decisions in a solitary carnivore. Behav Ecol 33, 137–146.

19. Silk, J.B. (2007). The adaptive value of sociality in mammalian groups. Philos Trans R Soc Lond B Biol Sci 362, 539–559.

20. Reser, J.E. (2014). Solitary mammals provide an animal model for autism spectrum disorders. J Comp Psychol 128, 99–113.

21. Alcock, J. (2013). Animal behavior: an evolutionary approach, 10th ed (Sinauer Associates).

22. Berry, R.J., and Bronson, F.H. (1992). Life history and bioeconomy of the house mouse. Biol Rev Camb Philos Soc 67, 519–550.

23. Luo, Z.X., Yuan, C.X., Meng, Q.J., and Ji, Q. (2011). A Jurassic eutherian mammal and divergence of marsupials and placentals. Nature 476, 442–445.

24. O’Leary, M.A., Bloch, J.I., Flynn, J.J., Gaudin, T.J., Giallombardo, A., Giannini, N.P., Goldberg, S.L., Kraatz, B.P., Luo, Z.X., Meng, J., et al. (2013). The placental mammal ancestor and the post-K-Pg radiation of placentals. Science 339, 662–667.

25. Porter, F.D., Drago, J., Xu, Y., Cheema, S.S., Wassif, C., Huang, S.P., Lee, E., Grinberg, A., Massalas, J.S., Bodine, D., et al. (1997). *Lhx2*, a LIM homeobox gene, is required for eye, forebrain, and definitive erythrocyte development. Development 124, 2935–2944.

26. Chou, S.J., and Tole, S. (2019). *Lhx2*, an evolutionarily conserved, multifunctional regulator of forebrain development. Brain Res 1705, 1–14.

27. Zhou, T., Zhu, H., Fan, Z., Wang, F., Chen, Y., Liang, H., Yang, Z., Zhang, L., Lin, L., Zhan, Y., et al. (2017). History of winning remodels thalamo-PFC circuit to reinforce social dominance. Science 357, 162–168.

28. Fan, Z., Zhu, H., Zhou, T., Wang, S., Wu, Y., and Hu, H. (2019). Using the tube test to measure social hierarchy in mice. Nat Protoc 14, 819–831.

29. Rein, B., Ma, K., and Yan, Z. (2020). A standardized social preference protocol for measuring social deficits in mouse models of autism. Nat Protoc 15, 3464–3477.

30. Jabarin, R., Netser, S., and Wagner, S. (2022). Beyond the three-chamber test: toward a multimodal and objective assessment of social behavior in rodents. Mol Autism 13, 41.

31. Shang, W., Xie, S., Feng, W., Li, Z., Jia, J., Cao, X., Shen, Y., Li, J., Shi, H., Gu, Y., et al. (2024). A non-image-forming visual circuit mediates the innate fear of heights in male mice. Nat Commun 15, 3746.

32. Moy, S.S., Nadler, J.J., Perez, A., Barbaro, R.P., Johns, J.M., Magnuson, T.R., Piven, J., and Crawley, J.N. (2004). Sociability and preference for social novelty in five inbred strains: an approach to assess autistic-like behavior in mice. Genes Brain Behav 3, 287–302.

33. Williamson, C.M., Lee, W., and Curley, J.P. (2016). Temporal dynamics of social hierarchy formation and maintenance in male mice. Anim Behav 115, 259–272.

34. Dehnen, T., Papageorgiou, D., Nyaguthii, B., Cherono, W., Penndorf, J., Boogert, N.J., and Farine, D.R. (2022). Costs dictate strategic investment in dominance interactions. Philos Trans R Soc Lond B Biol Sci 377, 20200447.

35. Clipperton-Allen, A.E., Almey, A., Melichercik, A., Allen, C.P., and Choleris, E. (2011). Effects of an estrogen receptor alpha agonist on agonistic behaviour in intact and gonadectomized male and female mice. Psychoneuroendocrinology 36, 981–995.

36. Barabas, A.J., Lucas, J.R., Erasmus, M.A., Cheng, H.W., and Gaskill, B.N. (2021). Who’s the boss? Assessing convergent validity of aggression based dominance measures in male laboratory mice, *Mus musculus*. Front Vet Sci 8, 695948.

37. Guillot, P.V., and Chapouthier, G. (1996). Intermale aggression and dark/light preference in ten inbred mouse strains. Behav Brain Res 77, 211–213.

38. Parmigiani, S., Palanza, P., Rogers, J., and Ferrari, P.F. (1999). Selection, evolution of behavior and animal models in behavioral neuroscience. Neurosci Biobehav Rev 23, 957–969.

39. Van Oortmerssen, G.A. (1971). Biological significance, genetics and evolutionary origin of variability in behaviour within and between inbred strains of mice (*Mus musculus*). A behaviour genetic study. Behaviour 38, 1–92.

40. LeClair, K.B., Chan, K.L., Kaster, M.P., Parise, L.F., Burnett, C.J., and Russo, S.J. (2021). Individual history of winning and hierarchy landscape influence stress susceptibility in mice. ELife 10, e71401.

41. Guo, S., Ji, W., Li, M., Chang, H., and Li, B. (2010). The mating system of the Sichuan snub-nosed monkey (*Rhinopithecus roxellana*). Am J Primatol 72, 25–32.

42. Huchard, E., English, S., Bell, M.B., Thavarajah, N., and Clutton-Brock, T. (2016). Competitive growth in a cooperative mammal. Nature 533, 532–534.

43. Vogt, C.C., Zipple, M.N., Sprockett, D.D., Miller, C.H., Hardy, S.X., Arthur, M.K., Greenstein, A.M., Colvin, M.S., Michel, L.M., Moeller, A.H., and Sheehan, M.J. (2024). Female behavior drives the formation of distinct social structures in C57BL/6J versus wild-derived outbred mice in field enclosures. BMC Biol 22, 35.

44. Lu, Y., Xu, X.R., Chen, B.Y., Jefferson, T.A., Fearnbach, H., and Yang, G. (2024). Spatiotemporal dynamics of the social structure of Indo-Pacific humpback dolphins (*Sousa chinensis*) in Xiamen waters from 2007 to 2019. Zool Res 45, 439–450.

45. Teichroeb, J.A., and Jack, K.M. (2017). Alpha male replacements in nonhuman primates: Variability in processes, outcomes, and terminology. Am J Primatol 79, e22674.

46. Chen, S., Xing, L., Xie, Z., Zhao, M., Yu, H., Gan, J., Zhao, H., Ma, Z., and Li, H. (2024). Single-cell transcriptomic reveals a cell atlas and diversity of chicken amygdala responded to social hierarchy. iScience 27, 109880.

47. Hiser, J., and Koenigs, M. (2018). The multifaceted role of the ventromedial prefrontal cortex in emotion, decision making, social cognition, and psychopathology. Biol Psychiatry 83, 638–647.

48. Abe, R., Okada, S., Nakayama, R., Ikegaya, Y.A.-O., and Sasaki, T. (2019). Social defeat stress causes selective attenuation of neuronal activity in the ventromedial prefrontal cortex. Sci Rep 9, 9447.

49. van der Kooij, M.A., and Sandi, C. (2015). The genetics of social hierarchies. Curr Opin Behav Sci 2, 52–57.

50. Lijam, N., Paylor, R., McDonald, M.P., Crawley, J.N., Deng, C.X., Herrup, K., Stevens, K.E., Maccaferri, G., McBain, C.J., Sussman, D.J., and Wynshaw-Boris, A. (1997). Social interaction and sensorimotor gating abnormalities in mice lacking *Dvl1*. Cell 90, 895–905.

51. Ma, M., Xiong, W., Hu, F., Deng, M.F., Huang, X., Chen, J.G., Man, H.Y., Lu, Y., Liu, D., and Zhu, L.Q. (2020). A novel pathway regulates social hierarchy via lncRNA AtLAS and postsynaptic synapsin IIb. Cell Res 30, 105–118.

